# A genomic region containing *RNF212* and *CPLX1* is associated with sexually-dimorphic recombination rate variation in Soay sheep (*Ovis aries*)

**DOI:** 10.1101/024869

**Authors:** Susan E. Johnston, Camillo Bérénos, Jon Slate, Josephine M. Pemberton

## Abstract

Meiotic recombination breaks down linkage disequilibrium and forms new haplotypes, meaning that it is an important driver of diversity in eukaryotic genomes. Understanding the causes of variation in recombination rate is important in interpreting and predicting evolutionary phenomena and for understanding the potential of a population to respond to selection. However, despite attention inmodel systems, there remains little data on how recombination rate varies at the individual level in natural populations. Here, we used extensive pedigree and high-density SNP information in a wild population of Soay sheep (*Ovis aries*) to investigate the genetic architecture of individual autosomal recombination rate. Individual rates were high relative to other mammal systems, and were higher in males than in females (autosomal map lengths of 3748 cM and 2860 cM, respectively). The heritability of autosomal recombination rate was low but significant in both sexes(*h^2^* = 0.16 & 0.12 in females and males, respectively). In females, 46.7% of the heritable variation was explained by a sub-telomeric region on chromosome 6; a genome-wide association study showed the strongest associations at the locus *RNF212*, with further associations observed at a nearby ~374kb region of complete linkage disequilibrium containing three additional candidate loci, *CPLX1*, *GAK* and *PCGF3*. A second region on chromosome 7 containing *REC8* and *RNF212B* explained 26.2% of the heritable variation in recombination rate in both sexes. Comparative analyses with 40 other sheep breeds showed that haplotypes associated with recombination rates are both old and globally distributed. Both regions have been implicated in rate variation in mice, cattle and humans, suggesting a common genetic architecture of recombination rate variation in mammals.

**AUTHOR SUMMARY:** Recombination offers an escape from genetic linkage by forming new combinations of alleles, increasing the potential for populations to respond to selection. Understanding the causes and consequences of individual recombination rates are important in studies of evolution and genetic improvement, yet little is known on how rates vary in natural systems. Using data from a wild population of Soay sheep, we show that individual recombination rate is heritable and differs between the sexes, with the majority of genetic variation in females explained by a genomic region containing the genes *RNF212* and *CPLX1*.

## INTRODUCTION

Recombination is a fundamental feature of sexual reproduction in nearly all multi-cellular organisms, and is an important driver of diversity because it rearranges existing allelic variation to create novel haplotypes. It can prevent the accumulation of deleterious mutations by uncoupling them from linked beneficial alleles (Muller 1964; Crow and Kimura 1965), and can lead to an increase in genetic variance for fitness, allowing populations to respond to selection at a faster rate (McPhee and Robertson 1970; Felsenstein 1974; Charlesworth and Barton 1996; Burt 2000): this is particularly true for small populations under strong selection, where beneficial and deleterious alleles are more likely to be linked (Hill-Robertson Interference), and their relative selective costs and benefits are likely to be stronger (Hill and Robertson 1966; Otto and Barton 2001). However, recombination may be associated with fitness costs; higher rates of crossing-over may increase deleterious mutations and chromosomal rearrangements (Inoue and Lupski 2002), or lead to the breakup of favorable combinations of alleles previously built up by selection, reducing the mean fitness ofsubsequent generations (Charlesworth and Barton 1996). Therefore, the relative costs and benefits of recombination are likely to vary within different contexts, leading to an expectation of variation in recombination rates within and between populations (Barton 1998; Burt 2000; Otto and Lenormand 2002).

Recent studies of model mammal systems have shown that recombination rates vary at an individual level, and that a significant proportion of variance is driven by heritable genetic effects (Kong *et al.* 2004; Dumont *et al.* 2009; Sandor *et al.* 2012). In cattle, humans and mice, the heritability of recombination rate is 0.22, 0.08—0.30 and 0.46. respectively, andgenome-wide association studies have repeatedly attributed some heritable variation to specific genetic variants, including *RNF212, CPLX1, REC8* and *PRDM9*, among others (Kong *et al.* 2008; Baudat *et al.* 2010; Sandor *et al.* 2012; Kong *et al.* 2014; Ma *et al.* 2015). The majority of these loci appear to influence crossover frequency, may have sex-specific or sexually-antagonistic effects on recombination rate (e.g. *RNF212* and *CPLX1* in humans and cattle; Kong *et al.* 2014; Ma *et al.* 2015) and may be dosage dependent (e.g. *RNF212* in mice; Reynolds *et al.* 2013). The locus *PRDM9* is associated with the positioning and proportion of crossovers that occur in mammalian recombination hotspots (i.e. regions of the genome with particularly high recombination rates; Baudat *et al.* 2010; Ma *et al.* 2015), although this locus is not functional in some mammal species, such as canids (Auton *et al.* 2013). These studies suggest that recombination rate has a relatively oligogenic architecture, and therefore has the potential to respond rapidly to selection over relatively short evolutionary timescales.

Such studies in model systems have provided key insights into the causes of recombination rate variation. However, with the exception of humans, studies have been limited to systems that are likely to have been subject to strong artificial selection in their recent history, a process that will favour alleles that increase recombination rate to overcome Hill-Robertson Interference (Hill and Robertson 1966; Otto and Barton 2001). Some experimental systems show increased recombination rates after strong selection on unrelated characters (Otto and Lenormand 2002), and recombination rates are higher in domesticated plants and animals compared to their progenitors (Burt and Bell 1987; Ross-Ibarra 2004; but see Muñoz-Fuentes *et al.* 2015). Therefore, artificial selection may result in different genetic architectures than exist in natural populations. Studies examining recombination rate in wild populations will allow dissection of genetic and environmental drivers of recombination rate, to determine if it is underpinned by similar or different genetic architectures, and ultimately will allow examination of the association between recombination rate and individual fitness, enabling understanding how this trait evolves in natural systems.

Here, we examine the genetic architecture of recombination rate variation in a wild mammal population. The Soay sheep (*Ovis aries*) is a Neolithic breed of domestic sheep that has lived unmanaged on the St Kilda archipelago, Scotland, U.K., since the Bronze age (Clutton-Brock *et al.* 2004). In this study, we integrate genomic and pedigree information to characterize autosomal cross-over positions in more than 3000 gametes in individuals from both sexes. Our objectives were as follows: 1) to determine the relative importance of common environment and other individual effects on recombination rates (e.g. age, sex, inbreeding coefficients); 2) to determine if individual recombination rates were heritable; 3) to identify specific genetic variants associated with recombination rate variation; and 4) to determine if the genetic architecture of recombination rate variation is similar to that observed in other mammal species.

## MATERIALS AND METHODS

### Study population and pedigree

Soay sheep living within the Village Bay area of the island of Hirta (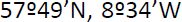) have beenstudied on an individual basis since 1985 (Clutton-Brock *et al.* 2004). All sheep are ear-tagged at first capture (including 95% of lambs born within the study area) and DNA samples for genetic analysis are routinely obtained from ear punches and/or blood sampling. All animal work was carried out according to UK Home Office procedures and was licensed under the UK Animals (Scientific Procedures) Act 1986 (License no. PPL60/4211). A Soay sheep pedigree has been constructed using 315 SNPs in low LD, and includes 5516 individuals with 4531 maternal and 4158 paternal links (Bérénos *et al.* 2014).

### SNP Dataset

A total of 5805 Soay sheep were genotyped at 51,135 single nucleotide polymorphisms (SNPs) on the Ovine SNP50 BeadChip using an Illumina Bead Array genotyping platform (Illumina Inc., San Diego, CA, USA; Kijas *et al.* 2009). Quality control on SNP data was carried out using the *check.marker* function in GenABEL v1.8-0 (Aulchenko *et al.* 2007) implemented in R v3.1.1, with the following thresholds: SNP minor allele frequency (MAF) > 0.01; individual SNP locus genotyping success > 0.95; individual sheep genotyping success > 0.99; and identity by state (IBS) with another individual < 0.90. Heterozygous genotypes at non-pseudoautosomal X-linked SNPs within males were scored as missing, accounting for 0.022% of genotypes. The genomic inbreeding coefficient (measure 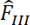 in Yang *et al.* 2011, hereafter 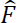), was calculated for each sheep in the software GCTA v1.24.3 (Yang *et al.* 2011), using information for all SNP loci passing quality control.

### Estimation of meiotic autosomal crossover count (ACC)

**Sub-pedigree construction**. To allow unbiased phasing of the SNP data, a standardized pedigree approach was used to identify cross-overs that had occurred within the gametes transferred from a focal individual to its offspring; hereafter, focal individual (FID) refers to the sheep in which meiosis took place. For each FID-offspring combination in the Soay sheep pedigree, a sub-pedigree was constructed to include both parents of the FID (Father and Mother) and the other parent of the offspring (Mate), where all five individuals had been genotyped (Figure 1). This sub-pedigree structure allowed phasing of SNPs within the FID, and thus the identification of autosomal cross-over events in the gamete transferred from the FID to the offspring (Figure 1). Sub-pedigrees were discarded from the analysis if they included the same individual twice (e.g. father-daughter matings; N = 13).

**Figure 1.**
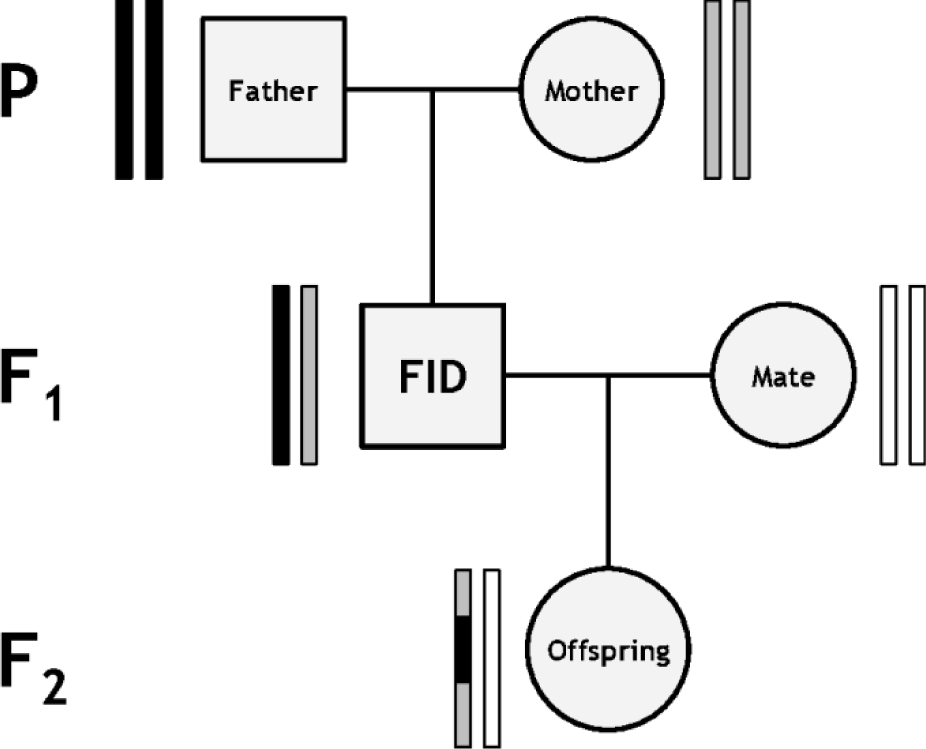
Diagram of the sub-pedigree structure used to infer crossover events. Rectangle pairs next to each individual represent chromatids, with black and grey shading indicating chromosome or chromosome sections of FID paternal and FID maternal origin, respectively. White shading indicates chromatids for which the origin of SNPs cannot be determined. Crossovers in the gamete transferred from the focal individual (FID) to its offspring (indicated by the grey arrow) can be distinguished at the points where origin of alleles origin flips from FID paternal to FID maternal and *vice versa*.

**Linkage map construction and chromosome phasing**. All analyses in this section were conducted using the software CRI-MAP v2.504a (Green *et al.* 1990). First, Mendelian incompatibilities in each sub-pedigree were identified using the *prepare* function; incompatible genotypes were removed from all affected individuals, and sub-pedigrees containing parent-offspring relationships with more than 0.1% mismatching loci were discarded. Second, sex-specific and sex-averaged linkage map positions (in Kosambi cM) were obtained using the *map* function, where SNPs were ordered relative to their estimated positions on the sheep genome assembly Oar_v3.1 (Genbank assembly ID: GCA_000298735.1; Jiang *et al.* 2014). SNP loci with a map distance of greater than 3 cM to each adjacent marker (10cM for the X chromosome, including PAR) were assumed to be incorrectly mapped and were removed from the analysis, with the *map* function rerun until all map distances were below this threshold; in total, 76 SNPs were assumed to be incorrectly mapped (these SNP IDs are included in archived data, see Data Availability). Third, the *chrompic* function was used to identify informative SNPs (i.e. those for which the grand parent of origin of theallele could be determined) on chromosomes transmitted from the FID to its offspring; crossovers were deemed to have occurred where there was a switch in the grandparental origin of a SNP allele (Figure 1).

**Quality control and crossover estimation in autosomes**. Errors in determining the grandparental origin of alleles can lead to false calling of double-crossovers (i.e. two adjacent crossovers occurring on the same chromatid) and in turn, an over-estimation of recombination rate. To reduce the likelihood of calling false crossover events, runs of grandparental-origin consisting of a single allele (i.e. resulting in a double crossover either side of a single SNP) were recoded as missing (N = 973 out of 38592 double crossovers, Figure S1). In remaining cases of double crossovers, the base pair distances between immediately adjacent SNPs spanning a double crossover were calculated (hereafter, “span distance” Figure S1). Informative SNPs that occurred within double-crossover segments with a log_10_ span distance lower than 2.5 standard deviations from the mean log_10_ span distance (equivalent to 9.7Mb) were also recoded as missing (N = 503 out of 37619 double crossovers, Figure S1). The autosomal crossover count (ACC), the number of informative SNPs and the informative length of the genome (i.e. the total distance between the first and last informative SNPs for all chromosomes) was then calculated for each FID. A simulation study was conducted to ensure that our approach accurately characterized ACC and reduced phasing errors. Autosomal meiotic crossovers were simulated given an identical pedigree structure and population allele frequencies (N_simulations_ = 100; see File S1 for detailed methods and results). Our approach was highly accurate in identifying the true ACC per simulation across all individuals and per individual across all simulations (adjusted R^2^ > 0.99), but indicated that accuracy was compromised in individuals with high values of 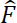. This is likely to be an artefact of long runs of homozygosity as a result of inbreeding, which may prevent detection of double crossovers or crossovers in sub-telomeric regions. To ensure accurate individual estimates of ACC, gametes with a correlation of adjusted R^2^ < 0.95 between simulated and detected crossovers in the simulation analysis were removed from the study (N = 8; File S1).

### Assessing variation in the recombination landscape

***Broad Scale Recombination Rate***. Relationships between chromosome length and linkage map length, and male and female linkage map length were analyzed using linear regressions in R v3.1.1. The relationship between chromosome length and chromosomal recombination rate (defined as cM length/Mb length) was modelled using a multiplicative inverse (1/*x*)regression in R v3.1.1.

***Fine Scale Recombination Rate***. The probability of crossing-over was calculated in1Mb windows across the genome using information from the male and female linkage maps, with each bin containing a mean of 15.6 SNPs (SD = 4.04). Briefly, the probability of crossing over within a bin was the sum of all recombination fractions, *r*, in that bin; in cases where an *r* value spanned a bin boundary, it was recalculated as *r* × N_boundary_/N_adjSNP_, where N_boundary_ was the number of bases to the bin boundary, and N_adjSNP_ was the number of bases to the closest SNP within the adjacent bin.

Variation in crossover probability relative to proximity to telomeric regions on each chromosome arm was examined using general linear models with a Gaussian error structure. The response variable was crossover-probability per bin; the fitted covariates were as follows: distance to the nearest telomere, defined as the best fit of either a linear (*x*), multiplicative inverse (1/*x*), quadratic (*x^2^* + *x*), cubic (*x^3^* + *x^2^* + *x*) or a log term (log_10_ *x*); sex, fitted as a main effect and as an interaction term with distance to the nearest telomere; number of SNPs within the bin; and GC content of the bin (%, obtained using sequence from Oar_v3.1, Jiang *et al.* 2014). The best model was identified using Akaike’s Information Criterion (Akaike 1974). An additional model was tested, using ratio of male to female crossover probability as the response variable, with the same fixed effect structure (omitting sex). In both models, the distance to the nearest telomere was limited to 60Mb, equivalent to half the length of the largest acrocentric chromosome (Chr 4). Initial models also included a term indicating if a centromere was present or absent on the 60Mb region, but this term was not significant in either model.

### Factors affecting autosomal recombination rate, including heritability and cross-sex genetic correlations

ACC was modelled as a trait of the FID. Phenotypic variance in ACC was partitioned using a restricted maximum likelihood (REML) Animal Model (Henderson 1975) implemented in ASReml-R (Butler *et al.* 2009) in R v3.1.1. To determine the proportion of phenotypic variance attributed to additive genetic effects (i.e. narrow-sense heritability, *h^2^*, hereafter heritability), a genomic relatedness matrix at all autosomal markers was constructed for all genotypedindividuals using GCTA v1.24.3 (Yang *et al.* 2011). The matrix was adjusted using the argument *—grm-adj 0*, which assumes that frequency spectra of genotyped and causal loci are similar; matrices with and without adjustment were highly correlated (R^2^ > 0.997) and variance components estimated from models with and without adjustment were highly similar, suggesting that adjusting for sampling error in this way did not introduce bias. Matrices were not pruned to remove related individuals (i.e. the *—grm-cutoff* option was not used) as there is substantial relatedness within this population, and models included common environment and parental effects, controlling for some consequences shared environment amongst relatives (see below). Trait variance was analyzed first with the following univariate model:

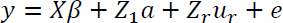

where *y* is a vector of the ACC per transferred gamete; *X* is an incidence matrix relating individual measures to a vector of fixed effects, beta; *Z_1_* and *Z_r_* are incidence matrices relating individual measures with additive genetic effects and random effects, respectively; *a* and *u_r_* are vectors of additive genetic effects from the genomic relatedness matrix and additional random effects, respectively; and *e* is a vector of residual effects. The heritability (*h^2^*) was calculated as the ratio of the additive genetic variance to the sum of the variance estimated for all random effects. Model structures were initially tested with a number of fixed effects, including sex, 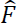 and FID age at the time of meiosis; random effects tested included: individual identity to account for repeated measures within the same FID (sometimes referred to as the permanent environment effect); maternal and paternal identity; and common environment effects of FID birth year and offspring birth year. Significance of fixed effects was determined using a Wald test, whereas significance of random effects was calculated using likelihood ratio tests (LRT) between models with and without the focal random effect. Only sex and additive genetic effects were significant in any model; however, 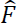 and individual identity were retained in all models to account for potential underestimation of ACC and the effects of pseudoreplication, respectively.

To investigate if the additive genetic variation underlying male and female ACC was associated with sex-specific variation in ACC, bivariate models were run. The additive genetic correlation *r_A_* was determined using the CORGH error structure function in ASReml-R, (correlation with heterogeneous variances) with *r_A_* set to be unconstrained; models fitted sex-specific inbreeding coefficients and individual identity effects. To test whether the genetic correlation was significantly different from 0 and 1, the unconstrained model was compared to models with *rA* fixed at a value of 0 or 0.999. Differences in additive genetic variance in males and females were tested by constraining both to be equal values using the CORGV error structure function in ASReml-R. Models were then compared using likelihood ratio tests with 1 degree of freedom.

### Genetic architecture of autosomal crossover count

**Genome-wide association study of variants controlling ACC**. Genome-wide association studies (GWAS) of autosomal recombination rates under different scenarios were conducted using ASReml-R(Butler *et al.* 2009) in R v3.1.1, fitting individual animal models for each SNP locus using the same model structure as above. SNP genotypes were fitted as a fixed effect with two or three levels. The GRM was replaced with a relatedness matrix based on pedigree information to speed up computation; the pedigree and genomic relatedness matrices have been shown to be highly correlated (Bérénos *et al.* 2014). Sex-specific models were also run. Association statistics were corrected for any population stratification not captured by the animal model by dividing them by the genomic control parameter, λ (Devlin *et al.* 1999), when λ > 1, which was calculated as the median Wald test 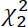 divided by the median 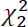 expected from a null distribution. The significance threshold after multiple testing was determined using a linkage disequilibrium-based method (outlined in Moskvina and Schmidt 2008) using a sliding window of 50 SNPs; the effective number of tests in the GWAS analysis was 22273.61, meaning the significance threshold for P after multiple testing at α = 0.05 was 2.245 χ 10^−6^. Although sex chromosome recombination rate was not included in the analysis, all GWAS included the X chromosome and SNP markers of unknown position (N=314). The proportion of phenotypic variance attributed to a given SNP was calculated using the following equation (Falconer and Mackay 1996):

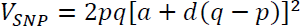

where *p* and *q* are the frequencies of alleles A and Bat the SNP locus, *a* is half the difference in the effect sizes estimated for the genotypes AA and BB, and *d* is the difference between *a* and the effect size estimated for genotype AB when fitted as a fixed effect in an animal model. The proportion of heritable variation attributed to the SNP was calculated as the ratio of V_SNP_ to the sum of V_SNP_ and the additive genetic variance estimated from a model excluding the SNP as a fixed effect. Standard errors of V_SNP_ were estimated using a delta method approach. Gene annotations in significant regions were obtained from Ensembl (gene build ID: Oar_v3.1.79; Cunningham *et al.* 2014). The position of a strong candidate locus, *RNF212* is not annotated on Oar_v3.1, but sequence alignment indicated that it is positioned at the sub-telomere of chromosome 6 (see File S2).

### Genome partitioning of genetic variance (regional heritability analysis)

Although a powerful tool to detect regions of the genome underlying heritable traits, the single locus approach of GWAS has reduced power to detect rare variants and variants with small effect sizes (Yang *et al.* 2011; Nagamine *et al.* 2012). One solution to this is to use a regional heritability approach that incorporates the effects of multiple haplotypes and determines the proportion of phenotypic variance explained by defined regions of the genome. The contribution of specific genomic regions to trait variation was determined by partitioning the additive genetic variance across all autosomes as follows (Nagamine *et al.* 2012):

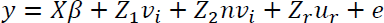

where *v* is the vector of additive genetic effects explained by an autosomal genomic region *i*, and *nv* is the vector of the additive genetic effects explained by all remaining autosomal markers outwith region *i*. Regional heritabilities were determined by constructing genomic relatedness matrices (GRMs) for regions of *i* of increasing resolution (whole chromosome partitioning, sliding windows of 150, 50 and 20 SNPs, corresponding to regions of 9.41 ± 1.42, 3.12 ± 0.60 and 1.21Mb mean ± 0.32 SD length, respectively) and fitting them in models with an additional GRM of all autosomal markers not present in region *i*; sliding windows overlapped by half of their length (i.e. 75, 25 and 10 SNPs, respectively). GRMs were constructed in the software GCTA v1.24.3 and were adjusted using the *–grm-adj 0* argument (see above; Yang *et al.* 2011). Adjusted and unadjusted matrices were highly correlated, but unadjusted matrices had higher incidences of negative pivots at the regional level. In cases where both models with adjusted and unadjusted matrices were used, there was little variation in estimated variance components, again suggesting that estimates were unbiased. The significance of additive genetic variance attributed to a genomic region *i* was tested by comparing models with and without the*Z_1_ v_i_* term using a likelihood ratio test; in cases where the heritability estimate was zero (i.e. estimated as "Boundary" by ASReml), significant model comparison tests were disregarded. A Bonferroni approach was used to account for multiple testing across the genome, by taking the number of tests and dividing by two to account for the overlap of the sliding windows (since each genomic region was modelled twice).

**Accounting for cis- and trans-genetic variants associated with recombination rate**. In the above analyses, we wished to separate potential associations with ACC due to cis-effects (i.e. genetic variants that are in linkage disequilibrium with polymorphic recombination hotspots) from those due to trans-effects (i.e. genetic variants in LD with genetic variants that affect recombination rate globally). By using the total ACC within a gamete, we incorporated both cis- and trans-effects into a single measure. To examine trans-effects only, we determined associations between each SNP and ACC minus crossovers that had occurred on the chromosome on which the SNP occurred e.g. for a SNP on chromosome 1, association was examined with ACC summed across chromosomes 2 to 26. We found that in this case, examining trans-variation (ACC minus focal chromosome) obtained similar results to cis- and trans-variation (ACC) for both regional heritability and genome-wide association analyses, leading to the same biological conclusions.

**Linkage disequilibrium and imputation of genotypes in significant regions**. A reference population of 189 sheep was selected and genotyped at 606,066 SNP loci on the Ovine Infinium^®^ HD SNP BeadChip for imputation of genotypes into individuals typed on the 50K chip. Briefly, the reference population was selected iteratively to maximize 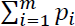 using the equation 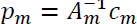, where *p* is a vector of the proportion of genetic variation in the population captured by *m* selected animals, *A_m_* is the corresponding subset of a pedigree relationship matrix and *c* is a vector of the mean relationship of the *m* selected animals (as outlined in Pausch *et al.* 2013 & Goddard and Hayes 2009). This approach should capture the maximum amount of genetic variation within the main population for the number of individuals in the reference population. SNP loci were retained if call rate was > 0.95 andMAF > 0.01 and individuals were retained if more than 95% of loci were genotyped. Linkage disequilibrium (LD) between loci was calculated using Spearman-Rank correlations (r^2^) inthe 188 individuals passing quality control.

Genotypes from the HD SNP chip were imputed to individuals typed on the SNP50 chip in the chromosome 6 region significantly associated with ACC, using pedigree information in the software MaCH v1.0.16 (Li *et al.* 2010). This region contained 10 SNPs from the Ovine SNP50 BeadChip and 116 additional independent SNPs on the HD SNP chip. As the software requires both parents to be known for each individual, cases where only one parent was known were scored as both parents missing.Genotypes were accepted when the dosage probability was between 0 and 0.25, 0.75 and 1.25, or 1.75and 2 (for alternate homozygote, heterozygote and homozygote, respectively). The accuracy of genotyping at each locus was tested using 10fold cross-validation within the reference population: genotypes were imputed for 10% of individuals randomly sampled from the reference population, using genotype data for the remaining 90%; this cross-validation was repeated 1000 times, to compare imputed genotypes with true genotypes. Cross-validation showed a relationship between number of missing genotypes and number of mismatching genotypes within individuals; therefore, individuals with < 0.99 imputed genotypes scored were removed from the analysis. Loci with < 0.95 of individuals typed were also discarded. Imputation accuracy was calculated for all loci as the proportion of imputed genotypes matching their true genotypes; all remaining loci had imputation accuracies >0.95.

### Haplotype sharing of associated regions with domesticated breeds

A recent study has shown that Soay sheep are likely to have experienced an introgression event with a more modern breed (the Old Scottish Shortwool, or Dunface breed, now extinct) approximately150 years ago (Feulner *et al.* 2013). Therefore, we wished to determine if alleles at the most highly associated imputed SNP, oar3_OAR6_116402578 (see Results), had recently introgressed into the population by examining haplotype sharing (HS) between Soay sheep and Boreray sheep, a cross between Dunface and Scottish Blackface sheep. We used data from the OvineSNP50 BeadChip data for Soays and a further 2709 individuals from 73 different sheep breeds (provided by the International Sheep Genomics Consortium, ISGC; see Kijas *et al.* 2012, Feulner *et al.* 2013 and Table S1). In both the Soay and non-Soay datasets of the Ovine SNP50 BeadChip, we extracted 58 SNPs corresponding to ~4Mb of the sub-telomeric region on chromosome 6 and phased them using Beagle v4.0 (Browning and Browning 2007). We identified core haplotypes of 6 SNP loci that tagged different alleles at oar3_OAR6_116402578. The length of HS between the core Soay haplotypes and non-Soay breeds was then calculated as follows: for each core haplotype *i* and each sheep breed *j*,any haplotypes containing *i* were extracted, and the distance from *i* to the first mismatching SNP downstream *i* was recorded. This was repeated for all pairwise comparisons of Soay and non-Soay haplotypes to determine a mean and standard deviation of HS between *i* and breed *j.*

### Data availability statement

The supplementary information contains information on additional analyses conducted and is referenced within the text. Table S1 contains the sex-averaged and sex-specific linkage map positions and genomic positions of SNP loci. Tables S3, S4 and S5 contain full detailed results and effect sizes of the regional heritability, genome-wide association and imputed association studies, respectively. The [Dryad] repository contains genomic data before and after quality control measures, pedigree information and sub-pedigree structures, autosomal genomic relatedness matrices, population-wide crossover probabilities and individual recombination rate results. All scripts for analysis are provided on a GitHub repository at https://github.com/susjoh/GENETICS_2015_185553.

## RESULTS

### Broad-scale variation in recombination landscape

We used information from 3330 sub-pedigrees and data from 39104 genome-wide SNPs typed on the Ovine SNP50 BeadChip (Kijas *et al.* 2009) to identify 98420 meiotic crossovers in gametes transferred from 813 unique focal individuals to 3330 offspring; this included 2134 offspring from 586 unique females and 1196 offspring from 227 unique males. A linkage map of all 26 autosomes had asex-averaged length of 3304 centiMorgans (cM), and sex-specific lengths of 3748 cM and 2860 cM in males and females, respectively, indicating strong male-biased recombination rates in this population (Male:Female linkage map lengths = 1.31; Figure S2, Table S2). There was a linear relationship between the length of autosomes in megabases (Mb) and linkage map lengths (cM; Adjusted R^2^ = 0.991, P < 0.001; Figure 2A). Chromosome-wide recombination rates (cM/Mb) were higher in smaller autosomes (fitted as multiplicative inverse function, adjusted R^2^ = 0.616, P < 0.001, Figure 2B), indicative of obligate crossing over. The degree of sex-differences in recombination rate based on autosome length in cM (i.e. differences in male and female recombination rate) was consistent across all autosomes (Adjusted R^2^ = 0.980, P < 0.001, Figure 2C).

**Figure 2.**
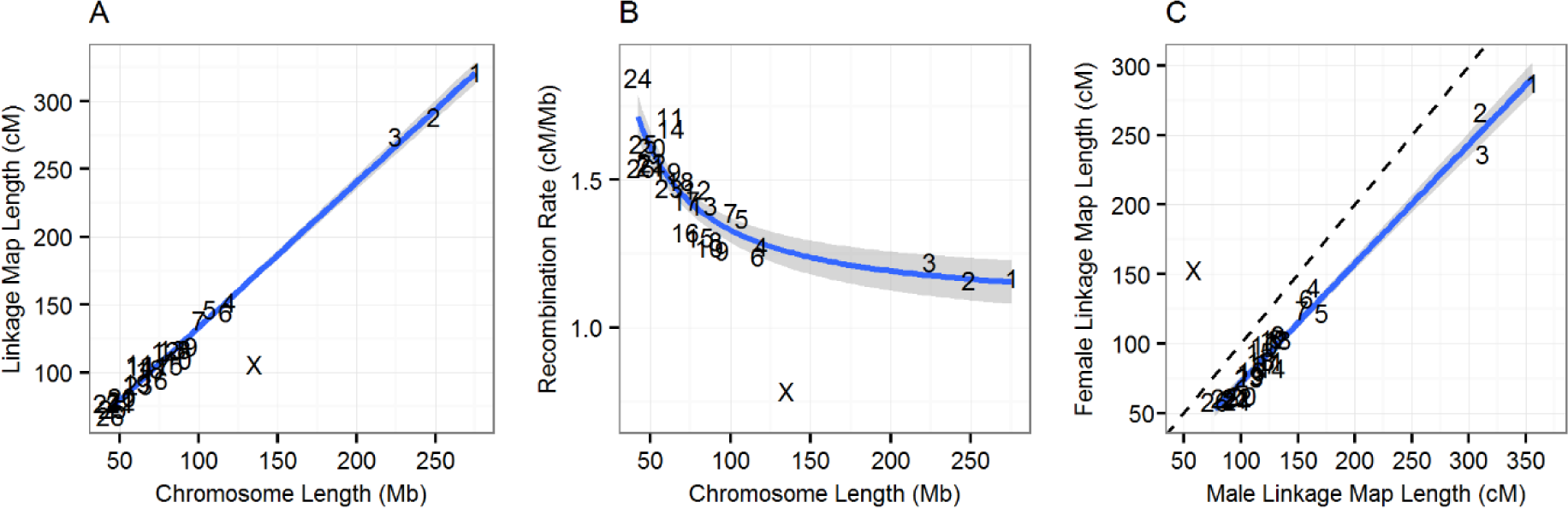
Broad scale variation in recombination rate. Relationships between: (A) sex-averaged linkage map length (cM) and physical chromosome length (Mb); (B) physical chromosome length (Mb) and recombination rate (cM/Mb); and (C) male and female linkage map lengths (cM). Points are chromosome numbers. Lines and the grey shaded areas indicate the regression slopes and standard errors, respectively, excluding the X chromosome. The dashed line in (C) indicates the where male and female linkage maps are of equal length. NB. The male linkage map length for the X chromosome is equivalent to the length of the pseudo-autosomal region.

### Fine-scale variation in recombination landscape

Finer-scale probabilities of crossing-over were calculated for 1Mb windows across the genome for each sex, using recombination fractions from their respective linkage maps. Crossover probability varied relative to proximity to telomeric regions, with a significant interaction between sex and distance to the nearest telomere fitted as a cubic polynomial function (Figure 3A). Males had significantly higher probabilities of crossing-over than females between distances of 0Mb to 18.11Mb from the nearest telomere (Figure 3B, Table S3). Increased crossover probabilities were associated with higher GC content (General linear model, P < 0.001; Table S3). Investigation of the relative distances between crossovers (in cases where two or more crossovers were observed on a single chromatid) indicated that there may be crossover interference within this population, with a median distance between double crossovers of 48Mb (Figure S1).

**Figure 3.**
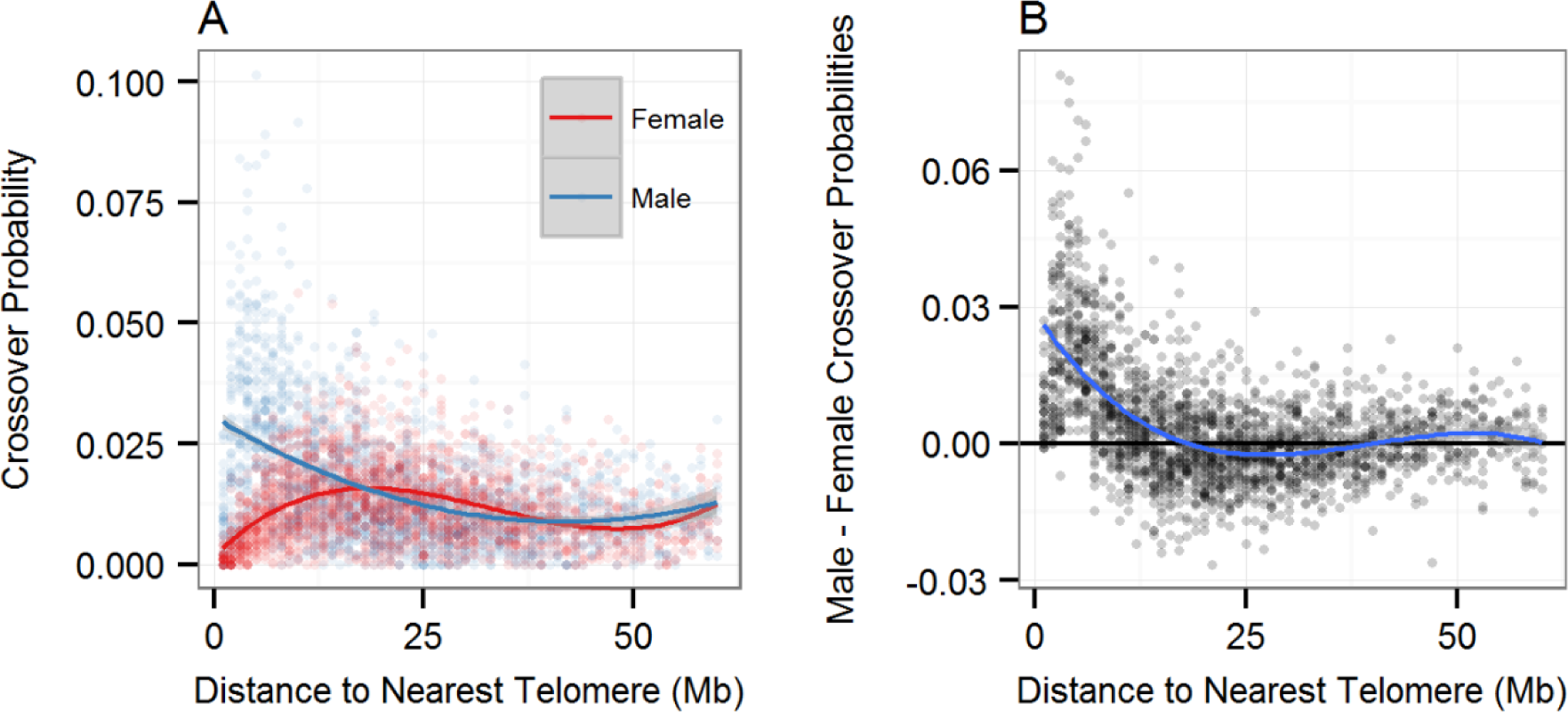
Variation in recombination rate relative to telomeric regions. Probability of crossing over relative to the nearest telomere (Mb) for (A) female and male linkage maps individually and (B) the difference between male and female crossover probabilities (male minus female). Data points are given for 1Mb windows. Lines indicate the function of best fit.

### Variation in individual recombination rate

Individual autosomal crossover count (ACC) was heritable (*h^2^* = 0.145, SE = 0.027), with the remainder of the phenotypic variance being explained by the residual error term (Table 1). ACC was significantly higher in males than in females, with 7.376 (SE = 0.263) more crossovers observed per gamete (Animal Model, Z = 28.02, P_Wald_ < 0.001). However, females had marginally higher additive genetic variance (P_LRT_ = 0.040) and higher residual variance (P_LRT_ = 1.11 × 10^−3^) in ACC than males (Table 1). There was no relationship between ACC and FID age, offspring sex, and the genomic inbreeding coefficient of the FID or offspring; furthermore, there was no variance in ACC explained by common environmental effects such as FID birth year, year of gamete transmission, or maternal/paternal identities ofthe FID (Animal Models, P > 0.05). A bivariate model of male and female ACC showed that the cross-sex additive genetic correlation (r_A_) was 0.826 (SE = 0.260); this correlation was significantly different from 0 (P_LRT_ = 1.14 × 10^−3^) but not different from1 (P_LRT_ = 0.551).

**TABLE 1.**
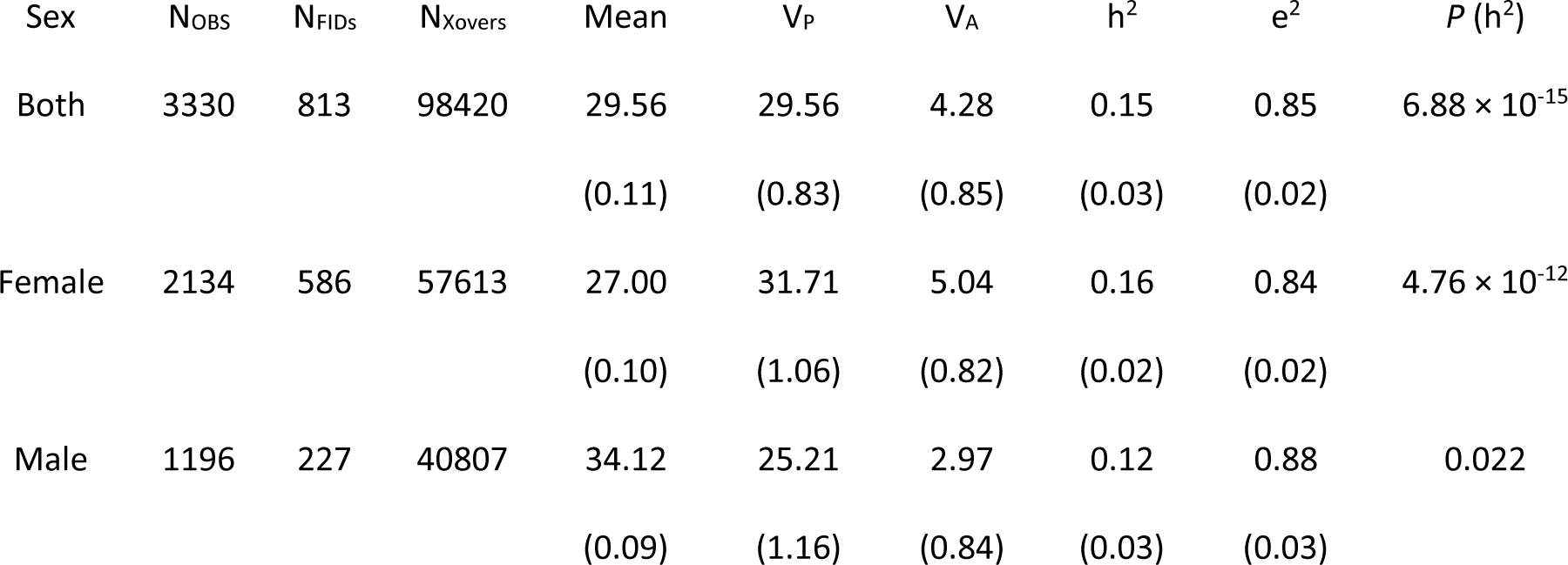
DATASET INFORMATION AND ANIMAL MODEL RESULTS OF AUTOSOMAL CROSSOVER COUNT (ACC) N_OBS_, N_FIDS_ and N_xovers_ indicate the number of ACC measures, the number of FIDs and the total number of crossovers observed, respectively. Mean is that of the raw data, and V_p_ and V_A_ are the phenotypic and additive genetic variances, respectively. The heritability h^2^ and residual effect e^2^ are the proportions of phenotype variance explained by the additive genetic and residual variances, respectively. *P*(h^2^) is the significance of the additive genetic effect (h^2^) as determined using a model comparison approach (see text). V_A_ and heritability were modelled using genomic relatedness. Figures in brackets are standard errors.

### Genetic architecture of recombination rate

**Genome-wide association study (GWAS)**. The most significant association between SNP genotype and ACC in both sexes was at s74824.1 in the sub-telomeric region of chromosome 6 (P = 2.92 × 10^−10^, Table 2). Sex-specific GWAS indicated that this SNP was highly associated with female ACC (P = 1.07 × 10^−11^), but was not associated with male ACC (P = 0.55; Table 2, Figure 4); the SNP had an additive effect on female ACC, with a difference of 3.37 (S.E. = 0.49) autosomal crossovers per gamete between homozygotes (Table 2). This SNP was the most distal typed on the chromosome from the Ovine SNP50 BeadChip at ~116.7Mb (Figure 4, Table 2), and corresponded to a genomic region containing ring finger protein 212 (*RNF212*) and complexin 1 (*CPLX1*), two loci that have previously been implicated in recombination rate variation in humans, cattle and mice (Kong *et al.* 2008, 2014; Sandor *et al.* 2012; Reynolds *et al.* 2013; Ma *et al.* 2015). A further SNP on an unmapped genomic scaffold (1.8kb, NCBI Accession: AMGL01122442.1) was also highly associated with female ACC (Figure 4). BLAST analysis indicated that the most likely genomic position of this SNP was at ~113.8Mb on chromosome 6, corresponding to the same sub-telomeric region.

**TABLE 2.**
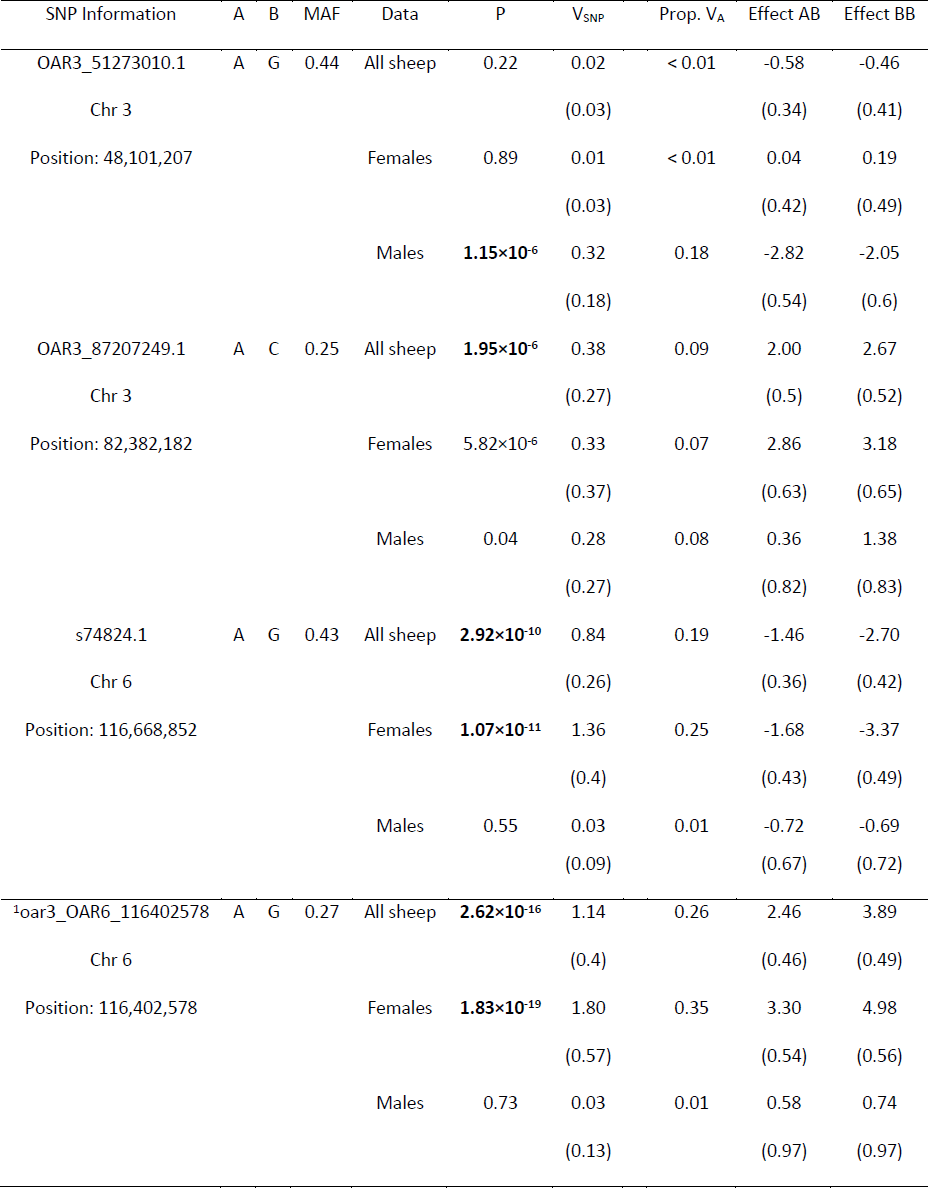
TOP HITS FROM GENOME-WIDE ASSOCIATION STUDIES OF ACC IN ALL SHEEP, FEMALES AND MALES. Results provided are from the Ovine SNP50 BeadChip and (below the line) the most highly associated imputed SNP from chromosome 6^1^. Additional loci that were significantly associated with ACC and in strong LD with these hits are not shown; full GWAS results are provided in Table S5 and S6. A and B indicate the reference alleles. P values are given for a Wald test of an animal model with SNP genotype fitted as a fixed effect; those in bold type were genome-wide significant. V_SNP_ is the variance attributed to the SNP and Prop.V_A_ is the proportion of the additive genetic variance explained by the SNP. Effect AB and BB are the effect sizes ofgenotypes AB and BB, respectively, relative to the model intercept at genotype AA. The number of unique individuals for all sheep, females and males are approximately N = 813,586 and 227, respectively. Numbers in brackets are standard errors.

**Figure 4.**
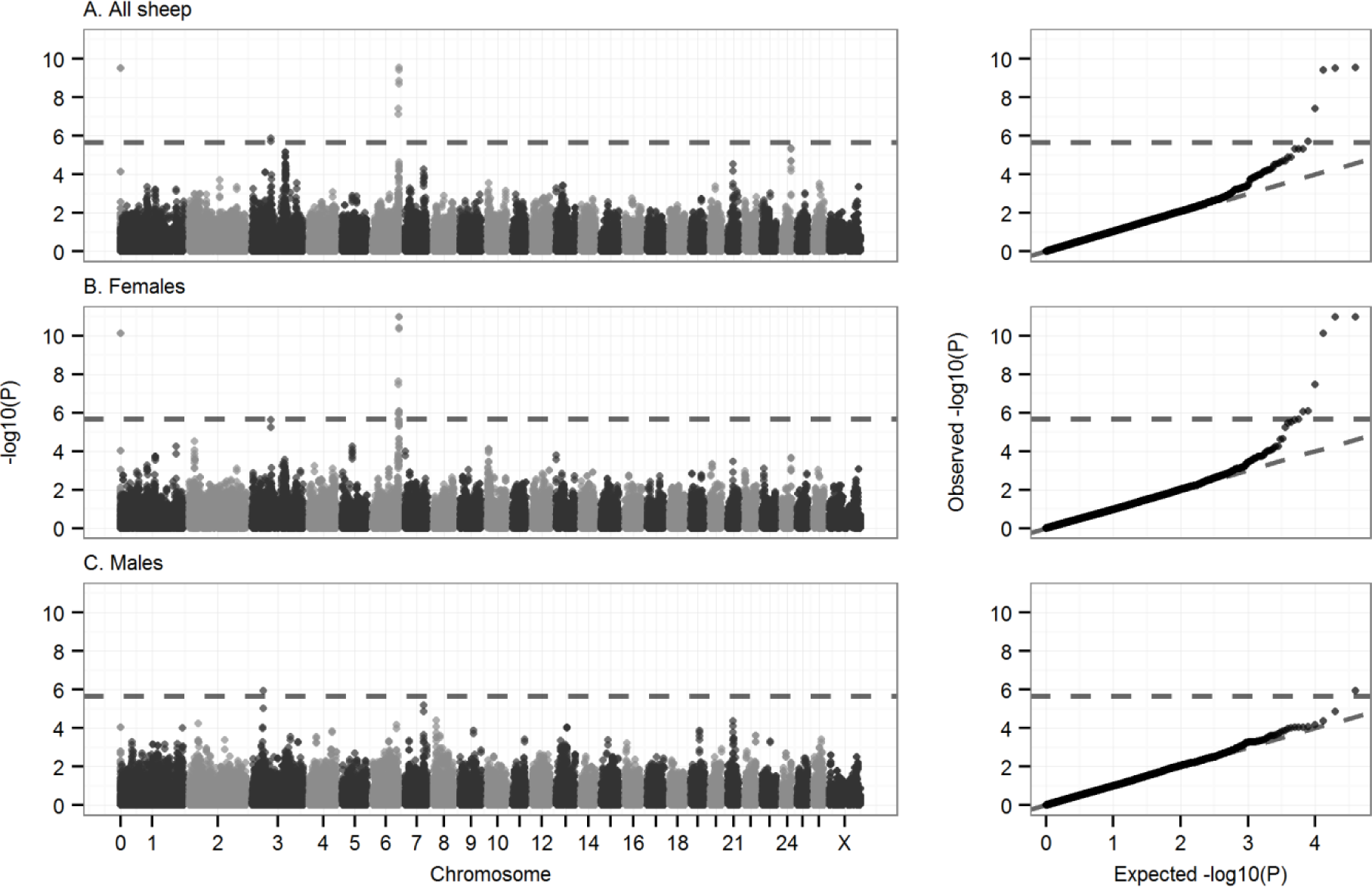
Genome-wide association of autosomal crossover count. Genome-wide association statistics in (A) all sheep, (B) females only and (C) males only. The dotted line indicates the threshold for statistical significance after multiple testing (equivalentto an experiment-wide threshold of P = 0.05). The left column shows association statistics relative to genomic position; points are color coded by chromosome. The right column shows the distribution of observed P values against those expected from a null distribution. Association statistics were not corrected using genomic control as λ was less than one for all GWAS (λ = 0.996, 0.933 and 0.900 for plots A, B and C, respectively). Underlying data on associations at the most highly associated SNPs, their genomic positions, and the sample sizes are given in Table S5. Thesignificant SNP in grey at position zero in (A) and (B) occurs on an unmapped contig that is likely to correspond to the distal region of chromosome 6 (see text).

Two further regions on chromosome 3 were associated with ACC using the GWAS approach. A single SNP, OAR3_51273010.1, was associated with ACC in males, but not in females, and had an approximately dominant effect on ACC (P = 1.15 × 10^−6^, Figure 4, Table 2); This SNP was 17.8kb fromthe 3~ UTR of leucine rich repeat transmembrane neuronal 4 (*LRRTM4*) in an otherwise gene poor region of the genome (i.e. the next protein coding regions are > 1Mb from this SNP in either direction). A second SNP on chromosome 3, OAR3_87207249.1, was associated with ACC in both sexes (P = 1.95x 10^−6^, Figure 4, Table 2). This SNP was 137kb from the 5΄ end of an orthologue of WD repeat domain 61 (*WDR61*) and 371kb from the 5΄ end of an orthologue of ribosomal protein L10 (*RPL10*). Full results of GWAS are provided in Table S4.

**Partitioning variance by genomic region**. The contribution of specific genomic regions toACC was determined by partitioning the additive genetic variance in sliding windows (Regional heritability analysis, Table S5). There was a strong sex-specific association of ACC in females withina sub-telomeric region on chromosome 6 (20 SNP sliding window; Figure 5B). This corresponded to a 1.46 Mb segment containing ~37 protein coding regions, including *RNF212* and *CPLX1.* The region explained 8.02% of the phenotypic variance (SE = 3.55%) and 46.7% of the additive genetic variance in females (PLRT = 9.78 χ 10^−14^), but did not contribute to phenotypic variation in males (0.312% of phenotypic variance, SE = 1.2%, P_LRT_ = 0.82; Figure 5C, Table S5). There was an additional significant association between ACC in both sexes and a region on chromosome 7, corresponding to a 1.09Mb segment containing ~50 protein coding regions, including *RNF212B* (a paralogue of *RNF212)* and meiotic recombination protein locus *REC8*(P_LRT_ = 3.31 × 10^−6^, Figure 5A, Table S5); this region had not shown any significant associations using the GWAS approach alone. The region explained 4.12% of phenotypic variance (SE = 2.3%) and 26.2% of the additive genetic variance in both sexes combined; however, in sex-specific models, the significant association with ACC did not remain after correction for multiple testing (Table S5). No association was observed in the regional heritability analysis with the two regions on chromosome 3 identified using the GWAS approach. Full results for the regional heritability analysis are provided in Table S5.

**Figure 5.**
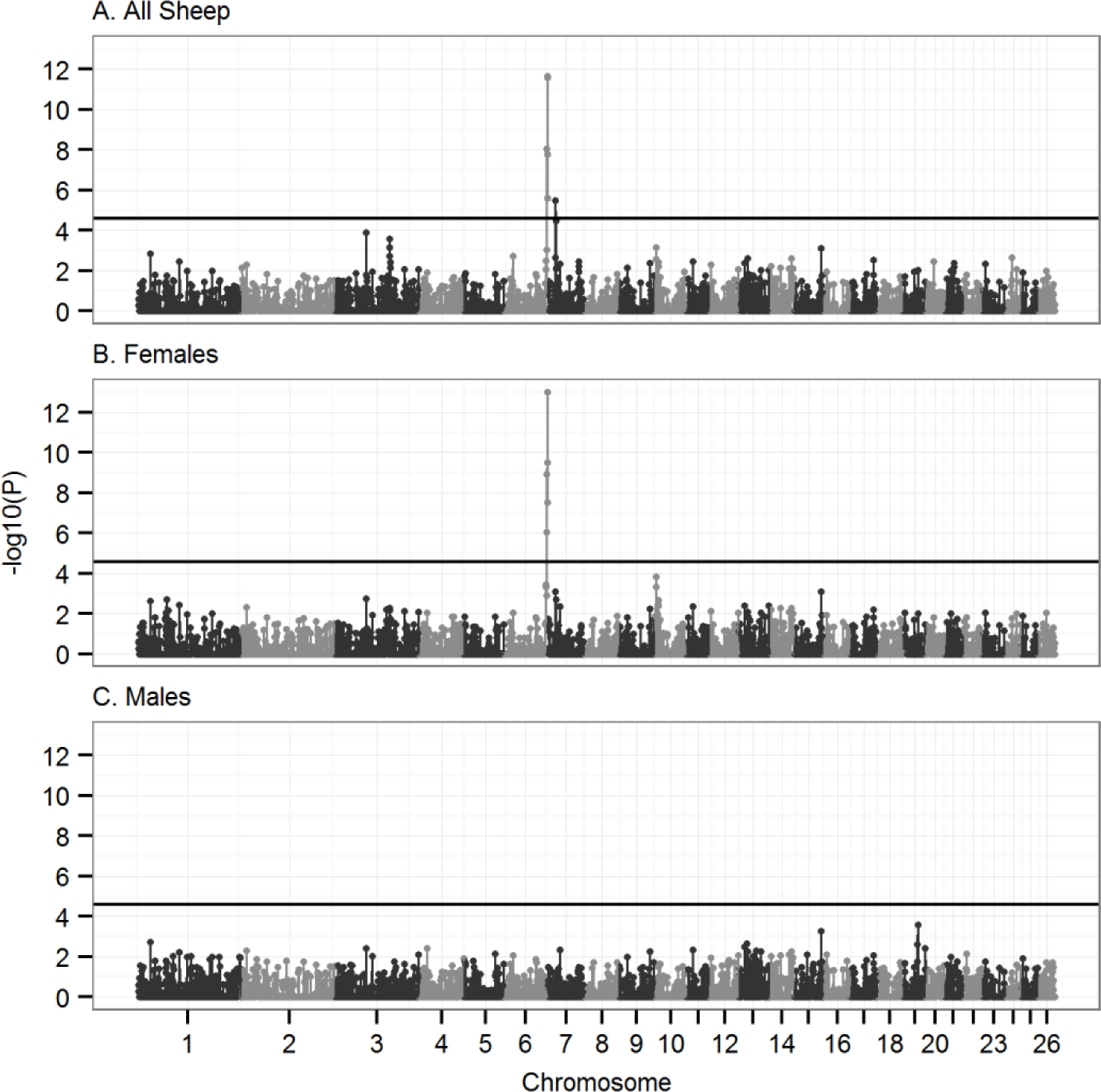
Regional heritability analysis of autosomal crossover count. Significance of association analysis in (A) all sheep, (B) females only and (C) males only. The results presented are from a sliding window of 20 SNPs across 26 autosomes, with an overlap of 10SNPs (see main text). Points represent the median base pair position of all SNPs within the sliding window. The solid black horizontal line is the significance threshold after multiple testing. Underlying data is provided in Table S4.

**Accounting for cis- and trans-genetic variants associated with recombination rate**. The results presented above were ACC incorporated both cis- and trans-effects on recombination rate. When repeated with trans-effects only, all variants associated with ACC in both the GWAS and regional heritability analyses remained significant (see Materials and Methods, Tables S4 & S5), meaning that they are likely to affect recombination rate globally (i.e. transacting effects), rather than being in LD with polymorphic recombination hotspots.

**Genotype imputation and association analysis at the sub-telomeric region of chromosome 6**. Genotyping of 187 sheep at a further 122 loci in the sub-telomeric region of chromosome 6 showed that this region has elevated levels of linkage disequilibrium, with the two most significant SNPs from the 50K chip tagging a haplotype block of ~374kB (r^2^ > 0.8; see File S3, Figure 6, Table S6). This block contained three candidate genes, complexin 1 (*CPLX1*), cyclin-G-associated kinase (*GAK*) and polycomb group ring finger 3 (*PCGF3*) and was 177kb away from the candidate locus *RNF212* (Kong *et al.* 2014). SNP genotypes were imputed for all individuals typed on the 50K chip at these 122 loci, and the association analysis was repeated. The most highly associated SNP (oar3_OAR6_116402578, P = 1.83 χ 10^−19^; Table 2, Figure 6) occurred within an intronic region of an uncharacterized protein orthologous to transmembrane emp24 protein transport domain containing (*TMED11*), 25.2kb from the putative location of *RNF212* and 13kb from the 3’ end of spondin 2 (*SPON2*). A bivariate animal model including an interaction term between ACC in each sex and the genotypeat oar3_OAR6_116402578 confirmed that this locus had an effect on female ACC only; this effect was additive, with a difference of 4.91 (S.E. = 0.203) autosomal crossovers per gamete between homozygotes (Figure 7, Tables 2 and S6). There was no difference in ACC between the three male genotypes. Full results for univariate models at imputed SNPs are given in Table S6.

**Figure 6.**
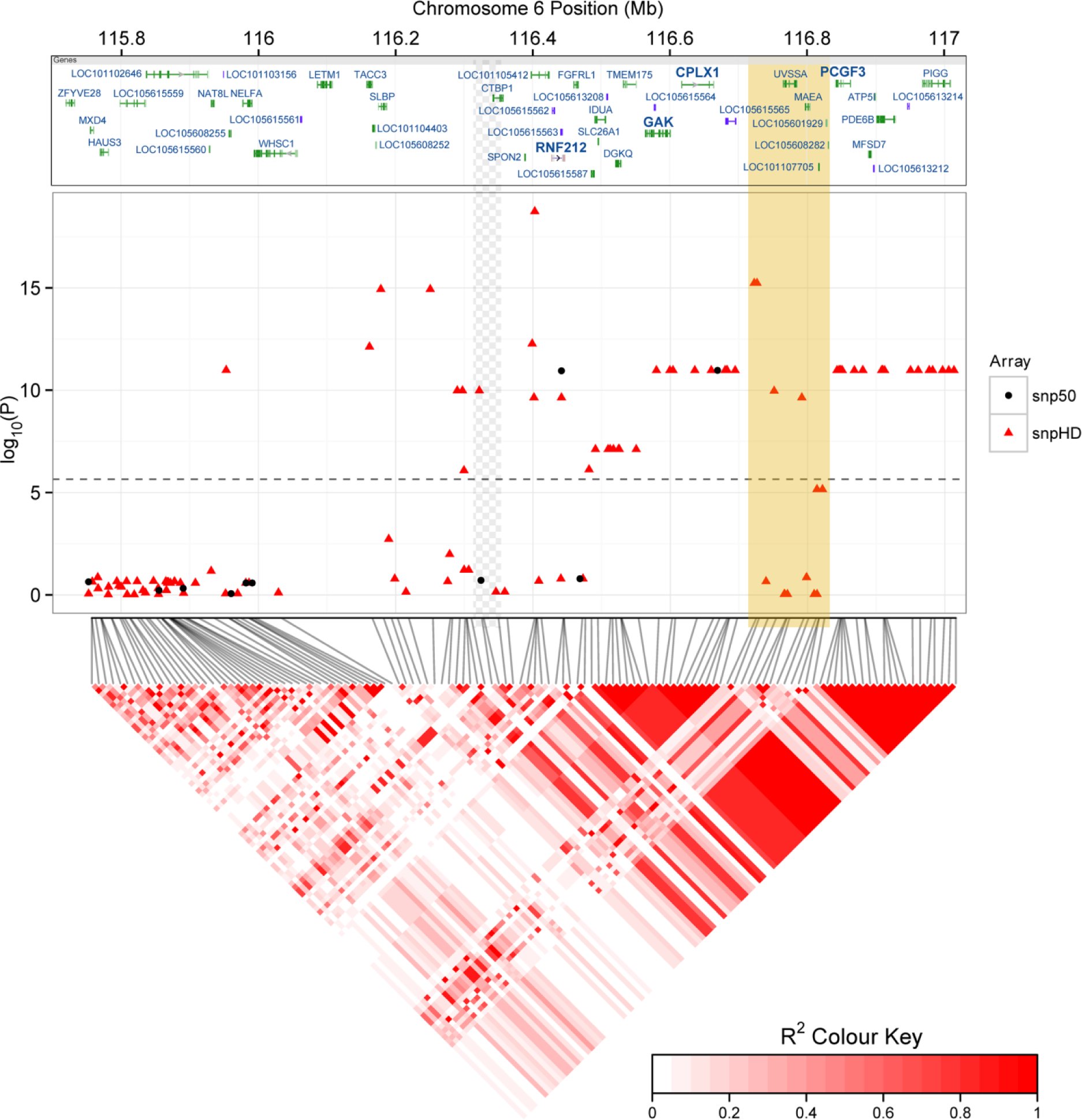
Associations at the sub-telomeric region of chromosome 6. Local associations of female ACC with Ovine SNP50 BeadChip SNPs (black circles, middle panel) and imputed genotypes from the Ovine HD SNP BeadChip (red triangles). The top panel indicates protein coding regions within this region as provided by the NCBI Graphical Sequence Viewer v3.8, with genes previously implicated in recombination or meiosis given in bold text (see Introduction and (Yokobayashi *et al.* 2013; Kong *et al.* 2014)). The dashed line in the middle panel indicates the significance threshold after multiple testing. The lower panel is a heatmap of linkage disequilibrium in this region calculated for the 188 individuals typed on the HD SNP chip using Spearman Rank correlation r^2^ created using the R library *LDheatmap*(Shin *et al.* 2006). The superimposed beige block indicates a region that is likely to be incorrectly assembled on the sheep genome assembly (Oar v3.1) based on sequence comparison with the homologous region of the cattle genome assembly (vUMD3.1; see File S3); its position is likely to fall within the region indicated by the grey chequered pattern to the left, leaving a large region of very high LD at the distal end of chromosome 6.

**Figure 7.**
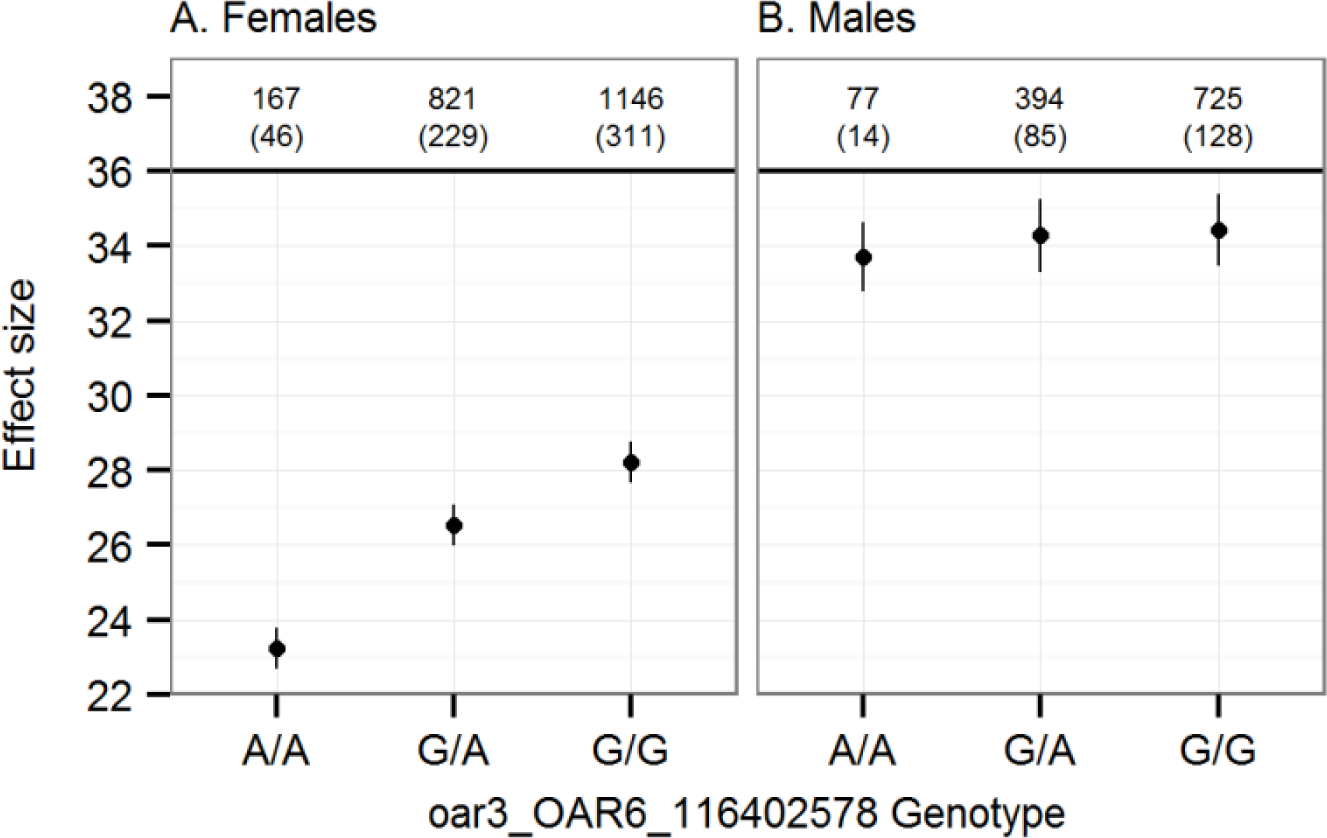
Effect sizes from a bivariate animal model of autosomal crossover count. Effect sizes are shown for (A) females and (B) males from a single bivariate model including oar3_OAR6_116402578 genotype as a fixed interaction term. Error bars are the standard error around the model intercept (genotype A/A) or the effect size relative to the intercept (genotypes G/A and G/G). Numbers above the points indicate the number of observations and the number of FIDs (in parentheses) for each genotype.

### Haplotype sharing of associated regions with domesticated breeds

Seven core haplotypes of six SNPs in length tagged different alleles at oar3_OAR6_116402578 at the sub-telomeric region of chromosome 6. Two were perfectly associated with the A allele at oar3_OAR6_116402578 (conferring reduced ACC) and five were perfectly associated with the G allele (conferring increased ACC; Table S8). The extent of HS between Soays and non-Soays was low, and there was no evidence of long-range HS between Soays and Boreray in comparison to other domesticated breeds (Figure S3). This test is not definitive due to the relatively small sample size of the Boreray sample (N = 20), meaning that it is possible that either allele occurs in Boreray sheep but has not been sampled. Nevertheless, low levels of haplotype sharing with other breeds throughout the sample suggest that allele at oar3_OAR6_116402578 have not been recently introduced to the Soay sheep population. For example, HS of core haplotypes with Boreray sheep for coat colour, coat pattern (Feulner *et al* 2013) and normal horn development (Johnston *et al.* 2013) extended to longer distances, of up to 5.7, 6.4 and 2.86Mb respectively. In contrast, the maximum HS observed here was 0.38Mb. A shorter haplotype may be expected, as the core haplotype occurs at the end of the chromosome, and so haplotype sharing is only calculated downstream of the core haplotype; however, this value is much lower than half that of previously identified introgressed haplotypes (Feulner *et al.* 2013). The three most common haplotypes, H2, H3 and H6 (for high, high and low ACC, respectively) are found in many other sheep breeds across the world (Figure S3), suggesting that both high and low ACC haplotypes are ancient across sheep breeds.

## DISCUSSION

In this study, we have shown that autosomal crossover count (ACC) is heritable in Soay sheep and that variation in female ACC is strongly influenced by a genomic region containing *RNF212* and *CPLX1*, loci that have previously been implicated in recombination rate variation in other species. The narrow sense heritability (*h^2^*) was 0.15 across both sexes, and was lower than estimates in some mammal species (*h^2^* = 0.22 and 0.46 in cattle and mice, respectively; Dumont *et al.* 2009; Sandor *et al.* 2012) and similar to recent estimates in humans (0.13 and 0.08 in females and males, respectively; Kong *et al.* 2014). ACCwas 1.3 times higher in males, but females had a higher proportion of heritable variation than males (*h^2^* = 0.16, compared to *h^2^* = 0.12). Here, we discuss the genetic architecture of the trait in more detail, the observation of sexual dimorphism and male-biased recombination rates, and how our findings inform the broader topic of understanding the genetic architecture of recombination rates in mammals.

### Genetic variants associated with individual recombination rate

The majority of variants associated with ACC in this study have previously been implicated in recombination rate variation in other mammal species, suggesting a shared genetic architecture across taxa. The strongest association was observed at the locus *RNF212*, occurring 88.4kB from a ~374kb block of high LD (r^2^ > 0.8) containing three further candidate loci, *CPLX1, GAK* and *PCGF3* (Figure 6). Both *RNF212* and *CPLX1* have been associated with recombination rate variation in mammals (Kong *et al.* 2008; Sandor *et al.* 2012; Reynolds *et al.* 2013; Ma *et al.* 2015) and mouse studies have established that the protein RNF212 is essential for the formation of crossover-specific complexes during meiosis, and that its effect is dosage-sensitive (Reynolds *et al.* 2013). We observed an additive effect of the *RNF212* region on female recombination rate (Figure 7), suggesting that dosage dependence could be a plausible mechanism driving rate differences in Soay sheep. GAK forms part of a complex with cyclin-G, a locus involved in meiotic recombination repair in *Drosophila* (Nagel *et al.* 2012), and PCGF3 forms part of a PRC1-like complex (polycomb repressive complex 1) which is involved in meiotic gene expression and the timing of meiotic prophase in female mice (Yokobayashi *et al.* 2013). High LD within this region meant that it was not possible to test the effects of these loci on recombination rate independently; however, the co-segregation of several loci affecting meiotic processes may merit further investigation to determine if recombination is suppressed in this region, and if this co-segregation is of adaptive significance.

Additional genomic regions associated with recombination rate included: two loci at 48.1Mb and 82.4Mb on chromosome 3 (identified using GWAS) with effects on males only and both sexes, respectively; and a 1.09Mb region of chromosome 7 affecting rates in both sexes (identified using regional heritability analysis). Although the chromosome 7 region was large and specific loci cannot be pinpointed, it contained *REC8*, the protein of which is required for the separation of sister chromatids and homologous chromosomes during meiosis (Parisi *et al.* 1999); and *RNF212B*,a paralogue of *RNF212*. The same region is also associated with recombination rate in cattle (Sandor *et al.* 2012). The chromosome 3 variants identified were novel to this study, and occurred in relatively gene poor regions of the genome (see above).

Although there are homologues of *PRDM9* on chromosomes 1, 5, 18 and X, it is not currently known whether any of these copies are functional in sheep. Here, we did not identify any association between recombination rate and any of these regions using either GWAS or regional heritability approaches. This may not be surprising, as this locus is primarily associated with recombination hotspot usage. Nevertheless, *PRDM9* has been associated with recombination rate in cattle and male humans (Kong *et al.* 2014; Ma *et al.* 2015) and is likely to be consequence of differences in the abundance of motifs recognized by the PRDM9 protein in hotspots rather than the locus itself affecting rate. In the current study, it was not possible to examine hotspot usage, as crossovers could only be resolved to a median interval of 800Mb. This is unlikely to be of the fine scale required to characterize hotspot variation, as they typically occur within 1-2kb intervals in mammals (Paigen and Petkov 2010). Further studies would require higher densities of markers to determine crossover positions at a greater resolution and to determine the functionality and/or relative importance of *PRDM9* within this system.

### Sexual dimorphism in genetic architecture of recombination rate

This study identified sexual dimorphism in the genetic architecture of recombination rate in Soay sheep. Using a classical quantitative genetic approach, the between-sex genetic correlation was not significantly different from 1, indicating that male and female recombination rate variation had a shared genetic basis – albeit with a relatively large error around this estimate. However, females had significantly higher additive genetic and residual variance in the trait in comparison to males, and GWAS and regional heritability showed that the *RNF212/CPLX1* region was associated with female recombination rate only. This is consistent with previous studies, where this region was associated with sexually dimorphic and sexually antagonistic variation in recombination rate in cattle and humans, respectively (Kong *et al.* 2014; Ma *et al.* 2015). Therefore, our findings suggest that variation in recombination rate has *some* degree of a shared and distinct genetic architecture between the sexes, which may be expected due to various similarities of this process of meiosis, but differences in its implementation within each sex (discussed further below). There were some differences in sample sizes between the sexes, with twice as many meioses characterized in females than males, so it could be argued that low sample sizes in males may have had less power to identify specific loci. Whilst possible, it is unlikely that the absence of associations between the *RNF212/CPLX1* region and male recombination rate is due to low power to detect the effect, as (a) models including both sexes showed reduced, rather than increased significance in this region; (b) bivariate models accounting for variation in *RNF212* as a fixed effect supported a sexually dimorphic genetic effect with a lower degree of error than the bivariate approach (Figure 7) and (c) repeating the association analysis at the most highly associated SNP using sampled datasets of identical size in males and females found consistently higher association at this locus in females (Figure S4).

### How much phenotypic variation in recombination rate is explained?

The approaches used in this study were successful in characterizing several regions of the genome contributing to the additive genetic variance in recombination rate. The regional heritability approach demonstrated some potential for characterizing variation from multiple alleles and/or haplotypes encompassing both common and rare variants that are in linkage disequilibrium with causal loci that were not detectable by GWAS alone (Nagamine *et al.* 2012). However, whilst some of the genetic contribution to phenotypic variance was explained by specific genomic regions,the overall heritability of recombination rate was low, and a substantial proportion of the heritable variation was of unknown architecture (i.e. “missing heritability” Manolio *et al.* 2009). In females, 64% of additive genetic variance was explained by the *RNF212/CLPX1* and *RNF212/REC8* regions combined, but this only accounted for 11% of the phenotypic variance (Table S9), leaving the remaining additive genetic and phenotypic variance unexplained. Our sample size is small relative to such studies in model systems, and there may have been reduced power to detect genetic variants, particularly in males which were under-represented inthe dataset. However, our finding of low heritability and unexplained additive genetic variance are consistent with recent results from Icelandic humans, where despite a higher sample size (N = 15,253 males and 20,674 females) and marker density (N = 30.3 × 10^6^), the fraction of phenotypic variance explained by specific loci remained small; identified variants including *RNF212* and *CPLX1* explained just 2.52 and 3.15% of male and female phenotypic variance, respectively, accounting for 29 and 24.8% of the additive genetic variance (Kong *et al.* 2014). Therefore, despite evidence of a conserved genetic architecture across mammal systems, there remains a very large proportion of both the additive genetic and phenotypic variance unexplained.

### Variation and sexual dimorphism in recombination landscape

Males had considerably higher recombination rates than in females, which was driven mainly by large differences in crossover frequencies in the sub-telomeric regions between 0 and 18.11Mb (Figure 3); recombination was reduced in males if the centromere was present in the sub-telomeric region (i.e. in autosomes 4 to 26, which are acrocentric), but unlike in cattle, was still significantly higher than that of females (Figure S5; Ma *et al.* 2015). Outside the sub-telomeric region, recombination rates were more similar between the sexes, with females showing slightly higher recombination rates between 18.1 and 40Mb from the telomere (Figure 3B). This observation of increased sub-telomeric recombination in males is consistent with studies in humans, cattle and mice (Kong *et al.* 2002; Shifman *et al.* 2006; Ma *et al.* 2015), although the magnitude of the difference is much greater in the Soay sheep population. Within females, the rate differences associated with different genotypes at *RNF212* were most clear in regions likely to be euchromatic, whereas there was no difference in rate in regions likely to be heterochromatic, such as the sub-telomeric and centromeric regions(Figure S6, Table S10).

### Why is recombination rate higher in males?

In placental mammals, females usually exhibit higher recombination rates than males (Lenormand and Dutheil 2005), and it has been postulated that this is a mechanism to avoid aneuploidy after long periods of meiotic arrest (Koehler *et al.* 1996; Morelli and Cohen 2005; Nagaoka *et al.* 2012). However, Soay sheep exhibited male-biased recombination rates to a greater degree than observed in any placental mammal to date (Male to Female linkage map ratio = 1.31). The biological significance of this remains unclear, although a number of mechanisms have been proposed to explain variation in sex differences more generally, including haploid selection (Lenormand and Dutheil 2005), meiotic drive (Brandvain and Coop 2012), sperm competition, sexual dimorphism and dispersal (Trivers 1988; Burt *et al.* 1991; Mank 2009). Nevertheless, testing these ideas has been limited by a paucity of empirical data.

One possible explanation for elevated recombination in males is that Soay sheep have a highly promiscuous mating system. Males have the largest testes to body size ratio within ruminants (Stevenson *et al.* 2004) and experiences high levels of sperm competition, with dominant rams becoming sperm-depleted towards the end of the annual rut (Preston *et al.* 2001). Increased recombination may allow more rapid sperm production through formation of meiotic bouquets (Tankimanova *et al.* 2004). Another argument made by Ma *et al.* (2015) to explain increased recombination in male cattle (M:F linkage map ratio = 1.1) is that stronger selection in males may have indirectly selected for higher recombination rates in bulls and may be a consequence of domestication (Burt and Bell 1987; Ross-Ibarra 2004). Soay sheep underwent some domestication before arriving on St Kilda, and have comparable levels of male-biased recombination to domestic sheep (M:F linkage map ratio = 1.19; Maddox *et al.* 2001). In contrast, wild bighorn sheep *(Ovis canadensis)* have not undergone domestication and have female-biased recombination rates (M:F linkage map ratio = 0.89; divergence ~2.8Mya; Poissant *et al.* 2010). However, low marker density (N = 232 microsatellites) in the bighorn sheep study may have failed to resolve crossovers in the sub-telomeric regions; furthermore, a recent study of chiasma count in wild progenitor vs. domestic mammal species found that recombination rates had not increased with domestication in sheep (Muñoz-Fuentes *et al.* 2015). In addition, wild cattle and sheep may have higher levels of sperm competition than other mammal species, supporting the former argument. Regardless, more empirical studies are required to elucidate the specific drivers of sex-differences in recombination rate at both a mechanistic and inter-specific level. Given our finding that the *RNF212/CPLX1* region is involved in the sex difference in Soay sheep, there is a compelling case for a role for this region in driving sex-differences in mammal systems over relatively short evolutionary timescales.

### Examining recombination rates in the wild

A principle motivation for the current study was to determine how recombination rate and its genetic architecture may vary relative to model species that have undergone strong selection in their recent history. We found that the heritability of recombination rate in Soay sheep was much lower than in cattle and mice, and was comparable with recent estimates in humans, which can also be considered a wild population (Kong *et al.* 2014). Despite these differences, variants affecting recombination rate on sheep chromosomes 6 and 7 have previously been associated with recombination rate in other mammalian populations (see above) and only the two relatively gene-poor regions identified on chromosome 3 are novel. Furthermore, examination of haplotypes around *RNF212* suggests that variation at the *RNF212/CPLX1* region affecting recombination rate has been segregating for a long time in sheep. Examining variation in the wild also allowed us to quantify the effects of the individual and environment on recombination rate; however, we found no effect of common environment effects (i.e. birth year and year of gamete transmission), individual age or inbreeding on recombination rate; rather, the majority of variation was attributed to residual effects. This contrasts with the observation in humans that recombination rates increase with age in females (Kong *et al.* 2004). Overall, our findings suggest a strong stochastic element driving recombination rates, with a small but significant heritable component that has a similar architecture to other mammal systems, regardless of their selective background. The Soay sheep system is one of the most comprehensive wild datasets in terms of genomic resources and sampling density, making it one of the most suitable in terms of quantifying and analyzing recombination rate variation in the wild. A future ambition is to investigate the fitness consequences of phenotypic variation in ACC and more specifically the relationship between variants identified in this study and individual life-history variation, to determine if the maintenance of genetic variation for recombination rates is due to selection, sexually antagonistic effects, or stochastic processes.

## Conclusions

In this study, we have shown that recombination rates in Soay sheep are heritable and have a sexually dimorphic genetic architecture. The variants identified have been implicated in recombination rates in other mammal species, indicating a conserved genetic basis across distantly related taxa. However, the proportion of phenotypic variation explained by identified variants was low; this was consistent with studies in humans and cattle, in which although genetic variants were identified, the majority of both additive genetic and phenotypic variance has remained unexplained. Similar studies in both mammalian and non-mammalian wild systems may provide a broader insight into the genetic and non-genetic drivers of recombination rate variation, as well its evolution in contemporary populations. However, the question remains as to whether variation in recombination rate is adaptive or merely a by-product of other biological processes. Overall, the approaches and findings presented here provide an important foundation for studies examining the evolution of recombination rates in contemporary natural populations.

## ACKNOWLEDGEMENTS

We thank Jill Pilkington, Ian Stevenson and all Soay sheep project members and volunteers for collection of data and samples. Discussions and comments from Jarrod Hadfield, Bill Hill, Craig Walling, Jisca Huisman, John Hickey and two anonymous reviewers greatly improved the analysis. James Kijas, Yu Jiang and Brian Dalrymple responded to numerous queries on the genome assembly, annotation and SNP genotyping. Philip Ellis prepared DNA samples, and Louise Evenden, Jude Gibson and Lee Murphy carried out SNP genotyping at the Wellcome Trust Clinical Research Facility Genetics Core, Edinburgh. This work has made extensive use of the resources provided by the Edinburgh Compute andData Facility (http://www.ecdf.ed.ac.uk/). Permission to work on St Kilda is granted by The National Trust for Scotland and Scottish Natural Heritage, and logistical support was provided by QinetiQ and Eurest. The Soay sheep project is supported by grants from the UK Natural Environment Research Council, with the SNP genotyping and research presented here supported by an ERC Advanced Grantto JMP.

